# Structural basis for rifamycin recognition and sequential substrate binding by the mycobacterial ADP-ribosyltransferase ARR

**DOI:** 10.64898/2026.05.04.722716

**Authors:** Lea C. von Soosten, Eike C. Schulz

## Abstract

The mycobacterial ADP-ribosyltransferase (ARR) mediates intrinsic antibiotic resistance by modifying the ansa-bridge of rifamycins with an ADP-ribose moiety derived from nicotinamide adenine dinucleotide (NAD^+^). Here, we elucidate the substrate binding mechanism of *M. smegmatis* ARR using X-ray crystallography and size-exclusion chromatography. We report the first apo structure of ARR, together with complexes involving rifampicin, rifabutin, and rifaximin. Biochemical data demonstrate an ordered sequential binding mechanism in which NAD^+^ requires prior rifamycin occupancy for recruitment to ARR. Structural comparison of apo ARR to rifamycin bound ARR, reveals that it undergoes significant conformational ordering upon rifamycin binding, characterized by a transition from highly flexible regions in the C-terminus and *α*2-helix to a more rigid state. Specifically, we identify a 2.9 Å shift in the loop between *β*-sheets 2 and 3, involving Phe39, which suggests a gating mechanism that organizes the NAD^+^ binding pocket. This experimental evidence is complemented by computational modelling of the NAD^+^ binding mode, which suggests a structurally plausible ternary arrangement. Together, these results establish that rifamycin binding is a prerequisite for NAD^+^ recruitment and provide a structural basis for the catalytic cycle of rifamycin ADP-ribosylation.

## 1 Introduction

The rising prevalence of infections caused by non-tuberculous mycobacteria (NTM) represents a formidable clinical challenge, largely due to the intrinsic resistance of these organisms to a broad spectrum of conventional antibiotics [1, 2]. Among the most essential therapeutic classes for treating mycobacterial infections are the rifamycins. These transcriptional inhibitors function by binding to the *β*-subunit (RpoB) of bacterial RNA polymerase, thereby obstructing the nascent RNA elongation process [3, 4]. Clinical applications of this class include rifampicin for tuberculosis therapy, rifabutin for the treatment of *Mycobacterium avium* complex, and rifaximin for gastrointestinal disorders [3].

While resistance to rifamycins is frequently driven by point mutations in *rpoB*, enzymatic inactivation via antibiotic modification has emerged as a significant mechanism of resistance [5]. A key mediator of this process is the rifampicin ADP-ribosyltransferase (ARR), an enzyme widely distributed across various bacterial species [6]. Unlike the more common role of ADP-ribosylation in protein post-translational modification and cellular regulation [7], ARR is unique in its ability to modify small-molecule antibiotics. First identified in the opportunistic pathogen *Mycobacterium smegmatis* [8], a frequently utilized model organism due to its high protein homology with pathogenic species such as *M. tuberculosis* and *M. abscessus* [9], ARR catalyzes the transfer of an ADP-ribose moiety from nicotinamide adenine dinucleotide (NAD^+^) to the ansa-bridge of the rifamycin substrate [10]. This modification renders the antibiotic incapable of binding to RNA polymerase, effectively neutralizing its bactericidal activity.

The proposed catalytic mechanism for ARR involves an *S*_*N*_ 1-type reaction, wherein NAD^+^ is cleaved to generate a reactive intermediate that is subsequently attacked by the C23-hydroxyl group of the rifamycin ansa-bridge [10]. Although the crystal structure of the ARR:rifampicin complex has been resolved [10, 11], critical questions regarding the enzyme’s kinetic and structural mechanism remain unanswered. Specifically, the precise binding order of the two substrates and the structural consequences of ligand recognition have not been elucidated. Furthermore, the lack of an apo-enzyme structure or complexes with other clinically relevant rifamycins has prevented a comprehensive understanding of the enzyme’s conformational landscape.

Here, we demonstrate that the substrates of *M*.*smegmatis* ARR bind in a strictly defined, ordered sequence. By combining size-exclusion chromatography with high-resolution X-ray crystallography, we present the first structural characterization of the ARR apo state and its complexes with rifampicin, rifabutin, and rifaximin. Together with computational modelling of a ternary ARR:rifamycin:NAD^+^ complex, our data reveal that rifamycin binding induces significant conformational changes that organize the NAD^+^ binding pocket, suggesting a structural gating mechanism essential for NAD^+^ binding.

## 2 Results

### Rifampicin binding is required for NAD^+^ recognition

To determine the substrate binding order of *M. smegmatis* ARR, we performed size-exclusion chromatography (SEC) experiments leveraging the distinct absorption maxima of the components (*λ*_*max*_: ARR, 280 nm; rifampicin, 495 nm; NAD^+^, 255 nm) and their characteristic elution volumes (Figure 1). Baseline elution profiles for the individual species were first established, showing peaks at ~0.5 CV for ARR, ~1.2 CV for rifampicin, and ~0.75 CV for NAD^+^ (Supplementary Figure 1a–c). We then evaluated the binding sequence through two complementary experiments:

**Figure 1:**
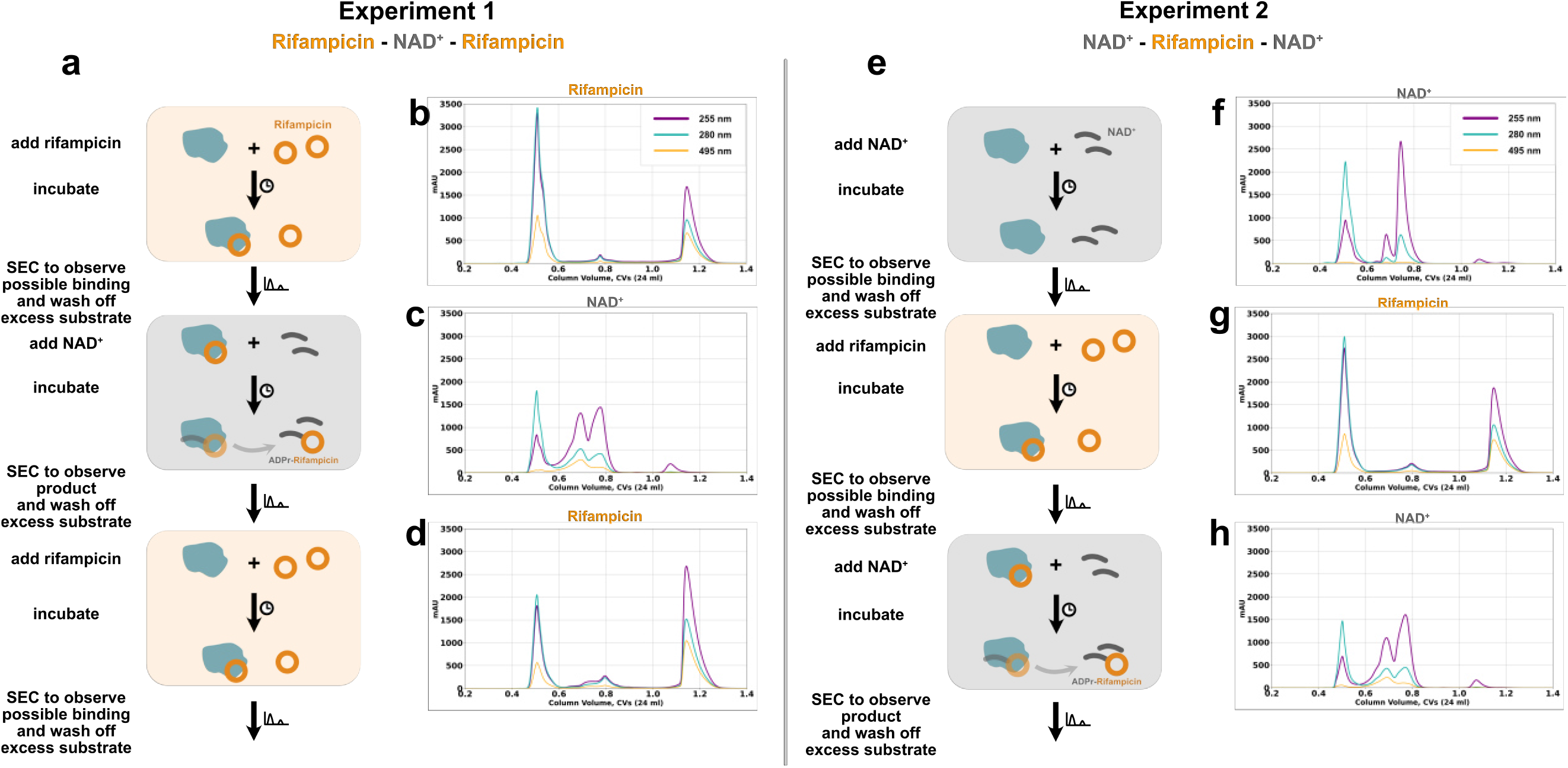
**a** and **e**: Schematic outline of the binding order experiment. Chromatograms showing that rifampicin binds to ARR, while NAD^+^ only binds to the ARR-rifampicin complex, followed by the reaction. **b** Rifampicin added to ARR, showing rifampicin binding to the protein. **c** NAD^+^ added to the ARR bound to rifampicin, the shift in absorption showing the product ADPr-rifampicin. **d** Rifampicin added to ARR, showing rifampicin binding to the protein. **f** NAD^+^ added to ARR, showing NAD^+^ to not bind to the protein. **g** Rifampicin added to ARR, showing rifampicin binding to the protein, but no product appearing on the chromatograms. **h** NAD^+^ added to ARR bound to rifampicin, the shift in absorption showing the product ADPr-rifampicin.

In Experiment 1, we tested whether rifampicin primes the enzyme for NAD^+^ recruitment. Incubating ARR with rifampicin resulted in a shift in the absorption spectrum and an elution peak at ~0.5 CV, confirming the formation of an ARR:rifampicin complex (Figure 1b). Subsequent addition of NAD^+^ to this preformed complex yielded a new elution peak at ~0.65 CV, alongside a minor rifampicin peak, indicative of a higher molecular weight species corresponding to the ADP-ribosylated product (Figure 1c). Furthermore, subsequent incubation with excess rifampicin produced peaks for both the ARR:rifampicin complex and unbound rifampicin, demonstrating that the enzyme remains capable of substrate binding following catalysis (Figure 1d).

In Experiment 2, we investigated whether NAD^+^ could bind to the apo enzyme independently. Incubation of ARR with NAD^+^ resulted in elution profiles for both species that remained unshifted, with an unaltered absorption ratio of 255:280 nm (Figure 1f), indicating no stable ARR:NAD^+^ complex was formed. While subsequent addition of rifampicin to the eluted ARR-peak allowed for the formation of an ARR:rifampicin complex (Figure 1g), ADP-ribosylation could only be triggered upon a secondary addition of NAD^+^, producing the characteristic product peak at ~0.65 CV (Figure 1h).

Taken together, these results unequivocally demonstrate an ordered, sequential binding mechanism in which the rifamycin must first occupy the active site, to render ARR competent for subsequent NAD^+^ recruitment. The observation that NAD^+^ fails to bind the apo enzyme (Figure 1f) hints upon rifampicin binding-induced conformational changes that organize the NAD^+^ binding pocket. To characterize these structural transitions, we next sought to obtain high-resolution structures of ARR in its apo state and in complex with various clinically relevant rifamycins.

### ARR displays high active site plasticity before substrate binding

To systematically investigate potential variations in rifamycin binding modes, we determined the crystal structures of *M*.*smegmatis* ARR in its apo state and as complexes with rifampicin, rifabutin, and rifaximin (Figure 2). Complexes were obtained via ligand soaking of existing crystals. All structures were solved by molecular replacement using the protein moiety of the previously reported structure (PDB ID 2HW2; [10]) as a search model. ARR crystallized in space group *C*2, with one molecule per asymmetric unit. The obtained structures were refined to resolutions ranging from 1.8 Åto 2.1 Å, with comprehensive refinement and data-collection statistics summarized in Table 2.

**Table 1.** Concentrations in the binding order experiment.

| Experiement | Run | ARR (mM) | Rifampicin (mM) | $\text{NAD}^+$ (mM) |
| --- | --- | --- | --- | --- |
| Reference | ARR | 0.25 |  |  |
|  | Rifampicin |  | 0.5 |  |
| | $\text{NAD}^+$ | | | 0.5 |
| Experiement 1 | 1 Rif | 0.25 | 0.5 |  |
| | 2 Rif - $\text{NAD}^+$ | collected | | 0.5 |
| | 3 Rif - $\text{NAD}^+$ - Rif | collected | 0.5 | |
| Experiement 2 | 1 $\text{NAD}^+$ | 0.25 | | 0.5 |
| | 2 $\text{NAD}^+$ - Rif | collected | 0.5 | |
| | 3 $\text{NAD}^+$ - Rif - $\text{NAD}^+$ | collected | | 0.5 |

**Table 2.**
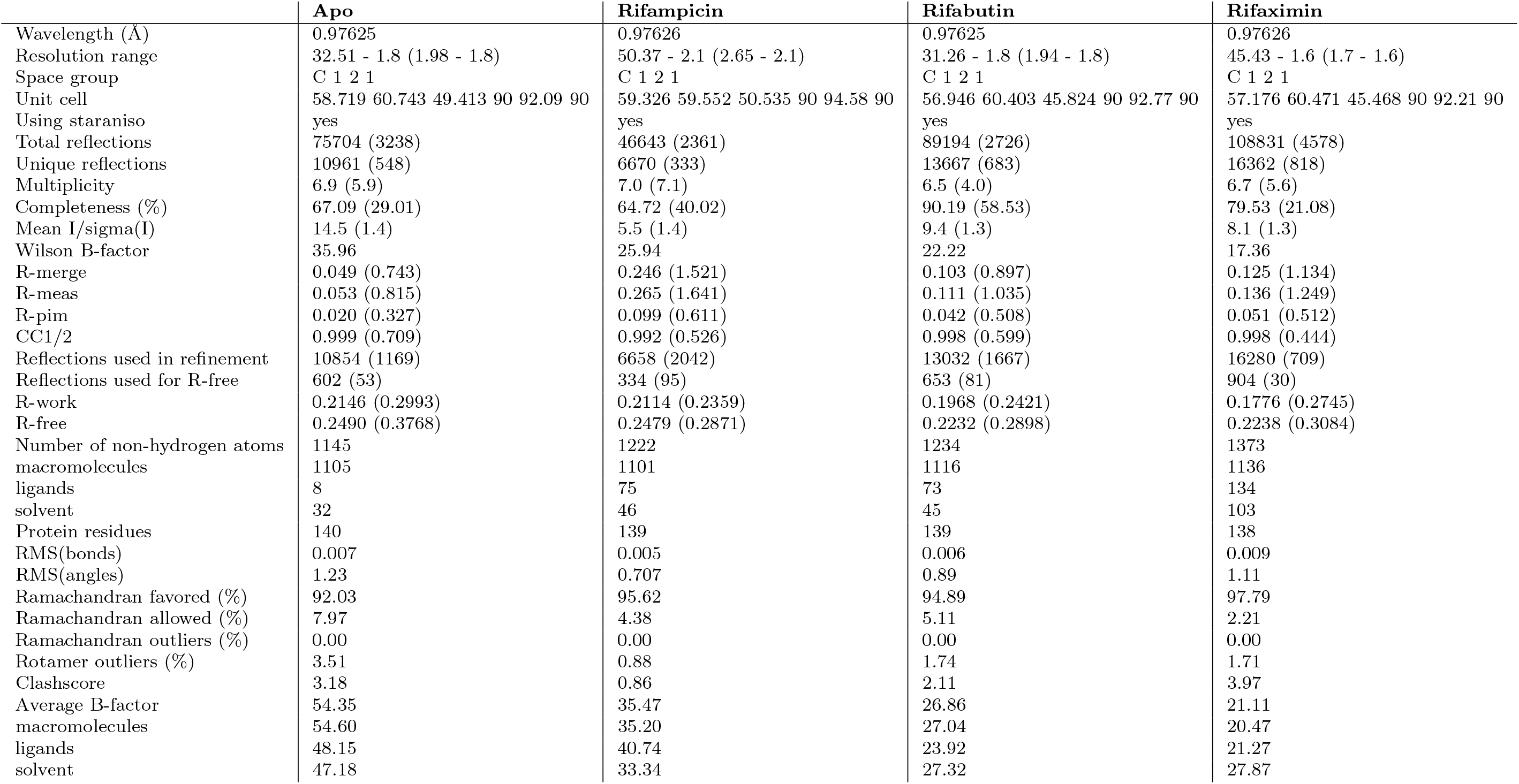
Data collection and refinement statistics.

**Figure 2:**
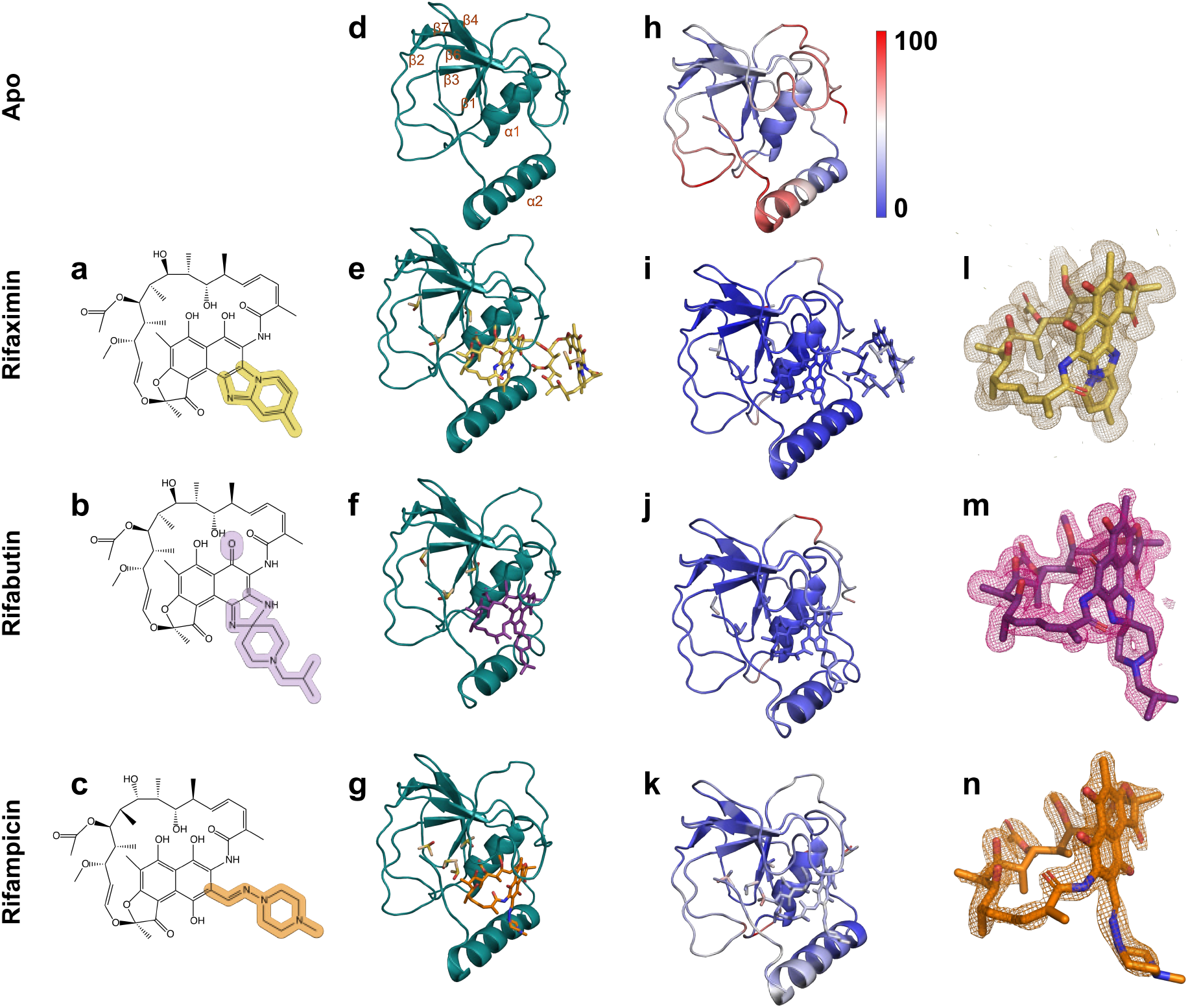
**a–c** Chemical structures of rifaximin, rifabutin and rifampicin. **d–g** Crystal structures of ARR as apo structure (**d**) and in complex with rifaximin (**e**), rifabutin (**f**) and rifampicin (**g**). **h–k** B-factor of the apo, rifaximin, rifabutin and rifampicin crystal structures. **l–m** Polder maps at 2.0 rmsd of rifaximin (**l**), rifabutin (**m**) and rifampicin (**n**).

The crystal structure of apo-ARR confirms that the enzyme adopts the single-domain fold characteristic of the rifamycin ADP-ribosyltransferase family [10]. The architecture is defined by a central *β*-sheet flanked by *α*-helices *α*1 and *α*2, which together delineate the deep cleft required for rifamycin recognition (Figure 2). Notably, the apo structure is characterized by significant conformational plasticity within the elements comprising the substrate-binding pocket. Compared to the more rigid ligand-bound states, we observed markedly increased B-factors in the C-terminus and the distal portion of the *α*2-helix (Figure 2h–k). Furthermore, both the loop containing LYS91, which acts as a lid over the binding cleft, and the flexible loop encompassing residue PHE39 between *β*-sheets 2 and 3, exhibit pronounced disorder. Collectively, these observations indicate that the active site is highly dynamic in the unliganded state, suggesting that rifamycin binding may facilitate a transition toward a more ordered and catalytically competent conformation.

### Rifamycin binding triggers ARR ordering

To characterize the impact of ligand binding on the observed conformational plasticity of ARR, we performed soaking experiments using apo crystals with rifampicin and the clinically relevant derivatives rifabutin and rifaximin (Figure 2a–c). In all cases, molecular replacement yielded robust models, confirmed by strong, well-defined positive difference electron density (DED) in the active site that is consistent with DED-omit maps (Figure 2l–m). Commonly all ARR:Rifamycin complexes display substantially lower B-factors in all areas lining the ligand-binding pocket, indicating that ligand binding triggers conformational ordering of ARR (Figure 2h–k).

The rifaximin complex exhibits a binding mode analogous to previously reported structures [10, 11]. Specifically, the O2 carbonyl of rifaximin is coordinated via hydrogen bonds to the backbone of ASN86 and LYS91, the latter of which resides on the flexible lid above the active site (Supplementary Figure 2a–b). The ansa-bridge is deeply embedded within the binding cleft, stabilized by a network of hydrophobic interactions and water bridges. We observed an additional rifaximin molecule coordinated at the interface between the ARR monomer and its symmetry mate. This second moiety appears to stabilize LYS91 through a *π*-cation interaction with the active-site rifaximin and a hydrogen bond with the symmetry-related molecule.

The rifabutin complex displays similar binding characteristics to rifaximin (Supplementary Figure 2c–d and 4), involving an analogous set of residues; however, it also engages THR59 through an additional hydrophobic interaction. Notably, LYS91 in the rifabutin structure is less rigidly coordinated, exhibiting higher B-factors compared to the rifaximin complex (Figure 2j). Further-more, variations in the binding orientations of LEU130 and LEU133 were observed relative to the rifaximin pose.

In contrast, the ARR:rifampicin complex exhibits a comparatively reduced contact surface area. While the overall orientation of rifampicin is consistent with previous reports [10, 11], we observed minor conformational discrepancies in the C15 carbonyl and the piperazine ring (Supplementary Figure 3). Most significantly, a major structural divergence was identified in the loop spanning *β*-sheets 2 and 3, encompassing residue PHE39. Our rifampicin structure reveals a 2.9 Åbackbone shift in this loop relative to both the rifabutin and rifaximin structures (Figure 3b). Interestingly, while the loops in the rifabutin and rifaximin complexes adopt a conformation consistent with the previously reported structure (PDB ID 2HW2), the loop in our rifampicin complex undergoes a distinct conformational shift towards the apo-like position. We speculate, that his variation could be influenced by the differing chemical environments in previous studies, such as the presence of the inhibitor Chr-16 (9IAF), glycylglycine (2HW2), or DMSO in our current structures, but further studies are required to provide more insight on this discrepancy. To further investigate whether these differences reflect distinct modes of substrate activation, we computationally modeled a ternary complex with the natural substrate NAD^+^.

**Figure 3:**
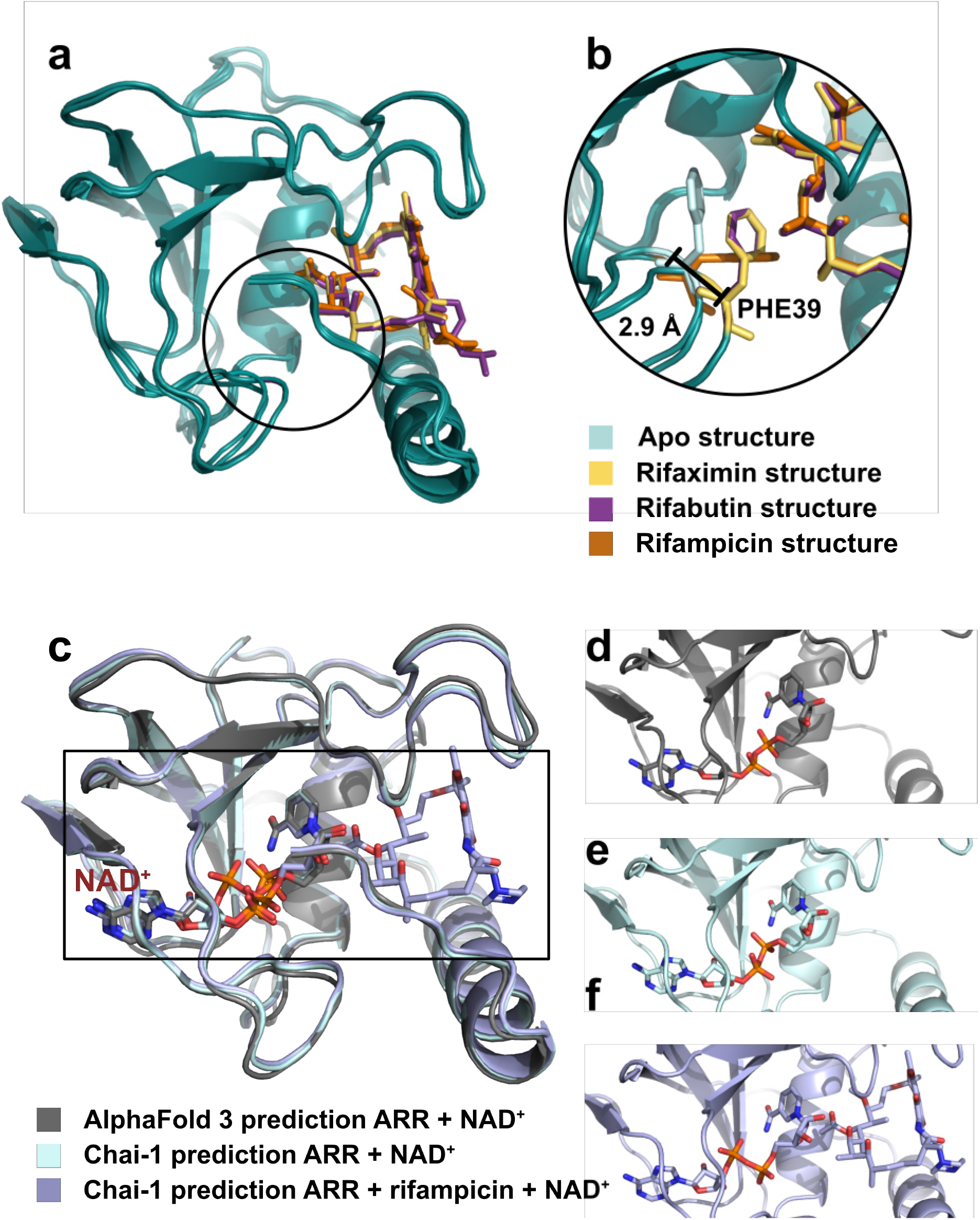
**a** and **b**: The loop between *β*-sheets 2 and 3 showing a backbone shift of 2.9 Åaround the PHE39 residue between the apo and rifampicin structures to the rifabutin and rifaximin structures. **c**: AlphaFold 3 [12] and Chai-1 [13] structure predictions. **c** Overlay of the three different predicted structures (**d** - **f**). **d** AlphaFold 3 prediction for ARR + NAD^+^. **e** Chai-1 prediction for ARR + NAD^+^. **f** Chai-1 prediction for ARR + rifampicin + NAD^+^.

### Computational modelling of the ARR:NAD^+^ binding landscape

To provide a structural rationale for the ordered substrate recruitment observed in our biochemical assays, we employed AlphaFold 3 [12] and Chai-1 [13] to predict the NAD^+^ binding mode within *M*.*smegmatis* ARR (Figure 3). Convergent predictions from both models identify a consistent, structurally plausible NAD^+^ binding site (Figure 3c), corroborating the location of the catalytic pocket. To specifically test whether rifampicin occupancy is required to organize the active site for NAD^+^ recruitment, we generated Chai-1 predictions for both a binary ARR:NAD^+^ complex and a ternary ARR:rifampicin:NAD^+^ complex. While the predicted binding pocket remains spatially conserved in both models, the NAD^+^ molecule adopts distinct conformational poses depending on the presence of rifampicin (Figure 3f). This computational insight supports our biochemical finding that NAD^+^ recruitment is coupled to rifampicin binding. Specifically, the computational data suggest that while a potential binding site exists in the apo state, the enzyme must undergo a conformational transition to stabilize a catalytically competent NAD^+^ pose. This transition appears to be driven by rifamycin binding to the deep cleft on the opposite side of ARR.

## 3 Discussion

In this study, we characterized the substrate binding mechanism of *M. smegmatis* ARR through a combination of biochemical and crystallographic approaches. Our size-exclusion chromatography experiments established an ordered sequential binding mechanism, demonstrating that NAD^+^ recruitment is strictly dependent on prior rifampicin occupancy.

This biochemical requirement is underpinned by the significant conformational plasticity observed in our structural data: while the apo-ARR structure is characterized by high B-factors and disorder in key regions, the enzyme undergoes substantial ordering upon rifamycin binding. The transition from a highly dynamic apo state to an ordered substrate-bound state suggests that ARR employs a conformational gating mechanism to regulate NAD^+^-binding. The loop encompassing residue PHE39, situated between the rifamycin and the predicted NAD^+^ binding sites, appears to play a central role in this process. The observed 2.9 Å backbone shift in this loop upon ligand binding suggests that it acts as a structural hinge or “gate” that must be repositioned to allow NAD^+^ access to the active site. This interpretation is further supported by the high degree of disorder observed in this region in the apo structure. Such an ordered recruitment mechanism ensures that the ADP-ribosylation reaction is only initiated when both substrates are correctly positioned, preventing wasteful NAD^+^ consumption. Whether this gating mechanism is a conserved feature among all rifamycin ADP-ribosyltransferases remains to be determined. The structural data reveal that despite the diverse chemical substituents found in different rifamycins (e.g., rifabutin, rifaximin), the core binding mode remains highly conserved. This conservation is reflected by consistent interactions with the naphthyl core and the ansa-bridge, providing a robust anchor within the binding cleft. Due to the large number of hydrophobic interactions to the ligands, ARR is able to maintain a broad substrate specificity, explaining its ability to inactivate both traditional and next-generation rifamycin derivatives.

While this substrate promiscuity is essential for the enzyme’s efficiency, it also underscores a significant challenge for clinical intervention: any new antibiotic designed to circumvent ARR must avoid these conserved interaction motifs to prevent rapid inactivation. Our findings expand the landscape of potential therapeutic strategies against ARR-mediated resistance. Based on our results we see two distinct yet plausible pharmacological approaches: (i) Competitive inhibition: Small molecules targeting the rifamycin binding site to physically block substrate access. (ii) Conformational or allosteric inhibition: Compounds designed to bind and stabilize the apo-like, disordered conformation of ARR, thereby preventing the structural transition required for NAD^+^ recruitment. The identification of this “gating” loop (PHE39) as a key regulatory element suggests that targeting these dynamic regions could provide a way to impede catalysis without necessarily competing directly with the rifamycin binding site. This is exemplified by the binding of the Chr-16 inhibitor in direct proximity to PHE39 (PDB-ID 9IAF), as recently demonstrated by Alaviuhkola et al. [11]. While our computational modeling provides a structurally plausible ternary arrangement, the precise orientation of NAD^+^ during the catalytic transition remains to be experimentally validated.

Future studies utilizing time-resolved serial crystallography (TR-SSX) will be essential to capture the real-time catalytic trajectory and provide a high-resolution view of the NAD^+^ binding and transfer process. Such advancements, combined with ongoing high-throughput screening efforts, will be critical in transitioning from structural characterization to the development of potent ARR inhibitors.

## 4 Materials and methods

### 4.1 Protein expression and purification

Codon-optimized *M. smegmatis* ARR (sequence provided in Supplementary Information) was cloned into a pETM-28a(+)CSSB vector (GenScript) containing an N-terminal His-SUMO tag with a SenP2 cleavage site. The construct was transformed into electrocompetent *E. coli* BL21gold cells and expressed in Terrific Broth (TB) supplemented with 100 *µ*g/ml kanamycin. Upon reaching an *OD*_600_ of *≈* 0.6, cultures were induced with 1 mM IPTG and incubated overnight at 21 °C. Cells were harvested and lysed via sonication in lysis buffer (50 mM HEPES pH 7.5, 500 mM NaCl, 2 mM MgCl_2_, 1 mM EDTA, 1 mg/ml lysozyme, 0.5 µg/ml DNAse, and protease inhibitors). The lysate was centrifuged and the supernatant filtered before it was subjected to immobilized metal affinity chromatography (IMAC) using a HisTrap column equilibrated 50 mM HEPES pH 7.5, 500 mM NaCl and 20 mM imidazole. The elution buffer contained 50 mM HEPES pH 7.5, 500 mM NaCl and 300 mM imidazole. The resulting fractions were pooled and dialysed overnight against 50 mM HEPES pH 7.5 and 300 mM NaCl (3.5 kDa MWCO) to facilitate SenP2-mediated SUMO-tag cleavage. To remove the cleaved tags and protease, a reverse IMAC step was performed using 50 mM HEPES pH 7.5, 300 mM NaCl and 20 mM imidazole for equilibration and 50 mM HEPES pH 7.5, 300 mM NaCl and 300 mM imidazole for the elution. The protein was concentrated (5 kDa MWCO) for the size-exclusion chromatography (SEC) using an SD75 HiLoad column equilibrated in 50 mM HEPES pH 7.5 and 300 mM NaCl. Final desalting and concentration were performed using a HiTrap Desalting column using 20 mM HEPES, pH 7.5. Purified ARR was flash frozen in liquid nitrogen and stored at −80 °C until further usage.

### 4.2 Binding order experiments

For the binding order experiment a SD75 increase column (Cytiva) was used for size exclusion chromatography. Before the experiment the column was washed with 1.2 CVs of water and then equilibrated with 1.2 CV 20 mM HEPES pH 7.5. The same buffer was used throughout the other steps. For the reference runs ARR, rifampicin and NAD^+^ were each applied to the column alone and eluted with 1.5 CVs buffer. It is important to note that rifampicin does elute much later (at ca. 1.2 CVs) than NAD^+^, even though it has a higher molecular weight.

#### Experiment-1

1 ml of *M. smegmatis* ARR (0.25 mM) was incubated at 30 °C for 60 minutes with twice the concentration of rifampicin (0.5 mM) added. The sample was then applied to the SEC column and eluted with 1.5 CVs of buffer. The fractions containing ARR were pooled and concentrated to 1 ml using a 5,000 MWCO centrifugal filter. Then NAD^+^ was added to a concentration of 0.5 mM. The sample was then again incubated for 60 minutes at 30 °C and run over the SEC column. Then the fractions containing ARR were collected, concentrated to 1 ml and rifampicin was added to a concentration of 0.5 mM. The sample was then applied to the SEC again and eluted with 1.4 CVs.

#### Experiment-2

1 ml of ARR (0.25 mM) was incubated at 30 °C for 60 minutes with twice the concentration of NAD^+^ (0.5 mM) added. The sample was then applied to the SEC column and eluted with 1.5 CVs of buffer. The fractions containing ARR were pooled and concentrated to 1 ml. Then rifampicin was added to a concentration of 0.5 mM. The sample was then again incubated for 60 minutes at 30 °C and run over the SEC column. The fractions containing ARR were collected, concentrated to 1 ml and NAD^+^ was added to a concentration of 0.5 mM. The sample was then applied to the SEC again and eluted with 1.5 CVs.

### 4.3 Protein crystallisation and complex formation

For protein crystallization ARR was concentrated to 30 mg/ml. ARR was crystallized via the sitting drop method with Swissci Maxi 48-Well Crystallization plates. The reservoir contained 200 µl of 75 % v/v MPD and 100 mM HEPES pH 7.5. The drops contained 1 µl of mother liquor and 2 µl of 30 mg/ml ARR in 20 mM HEPES pH 7.5. Crystals started forming after 1-2 days at 21 °C. For the soaking of the ARR crystals with the antibiotics rifampicin, rifabutin and rifaximin the antibiotics were each dissolved at 60 mM in 100 % DMSO. 1 µl of the solution was added directly to the crystallisation drop and incubated for 60 minutes. The soaked crystals were then fished and flash frozen in liquid nitrogen and stored therein prior to data-collection.

### 4.4 Data collection and structure determination

All datasets were collected at the P14 beamline at PETRA-III (EMBL Unit Hamburg, Germany) using an EIGER2 CdTe 16M detector (Dectris, Baden-Daetwill, Switzerland) at an energy of 12.7 keV (0.9763 Å). For every dataset 3600 images were collected. For the apo structure data collection the flux had 3.81 *·* 10^11^ ph s^*−*1^, for the rifampicin structure flux 9.71 *·* 10^10^ ph s^*−*1^, for the rifabutin structure 3.14 *·* 10^10^ ph s^*−*1^ and for rifaximin at 1.37 *·* 10^11^ ph s^*−*1^. The transmission of the beam was always set to 20 %. Diffraction data were processed in autoPROC v1.0.5 including Staraniso [14, 15], via XDS vJun30 2023 [16] and scaled in AIMLESS v0.8.2[17]. The structures were solved using molecular replacement in PHASER [18] using the structure PDB-ID 2HW2 [10] as a search model for ARR. For the structure refinement iterative cycles of phenix.refine (Phenix 2.0-5936, [19] were used and COOT-v0.9.8.95 [20] for the iterative cycles of model building. Data collection and refinement statistics are summarised in Table 2.

### 4.5 Data visualization

Molecular graphics were generated using PyMol [21], PLIP [22], and ChemDraw (Revvity Signals Software, Inc.).

### 4.6 Computational protein-ligand modeling

To investigate potential ternary complex arrangements, structural predictions were generated using AlphaFold 3 [12] and Chai-1 [13]. For all simulations, the protein sequence employed for crystallization was used as input.

#### AlphaFold 3

Predictions of the ARR–NAD^+^ complex were generated using one NAD^+^ molecule and an automatically generated seed (1978369222). For all comparative analyses, model 0 was selected based on the highest predicted model confidence metrics.

#### Chai-1

Predictions for both the binary ARR–NAD^+^ and ternary ARR:rifampicin:NAD^+^ complexes were performed using the same protein sequence. Multiple Sequence Alignment (MSA) generation was conducted via MMseqs2 with no templates or restraints applied. Ligands were provided as SMILES strings:

Rifampicin:

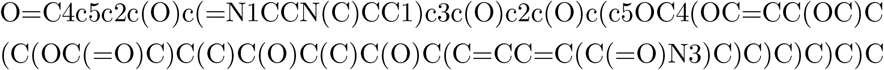

NAD^+^:

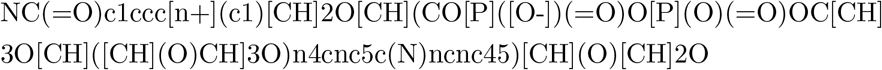

The confidence metrics of all predictions are shown in Supplementary Figure 5.

## Acknowledgements

All data was collected at beamline P14 operated by EMBL Hamburg at the PETRA-III storage ring (DESY, Hamburg, Germany).

## Author contributions

E.C.S. designed the experiments; LvS. purified and crystallized the protein, and carried out the ligand binding experiments, performed the data collection, processed and analysed the diffraction data, and refined the structures. L.vS. and E.C.S. wrote the manuscript; All authors discussed and corrected the manuscript.

## A Supplementary Figures

**Supplementary Figure 1:**
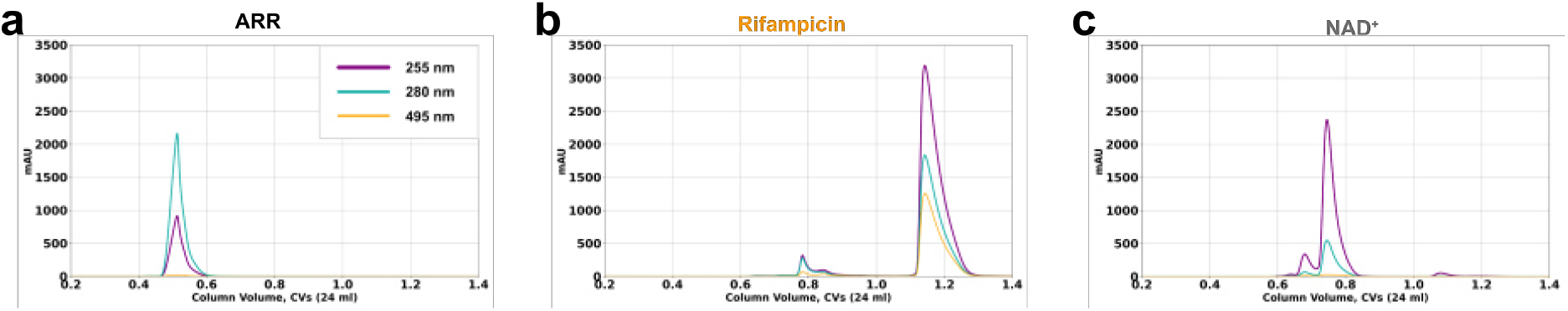
Size-exclusion chromatograms showing **a** ARR only, **b** rifampicin only, **c** NAD^+^ only.

**Supplementary Figure 2:**
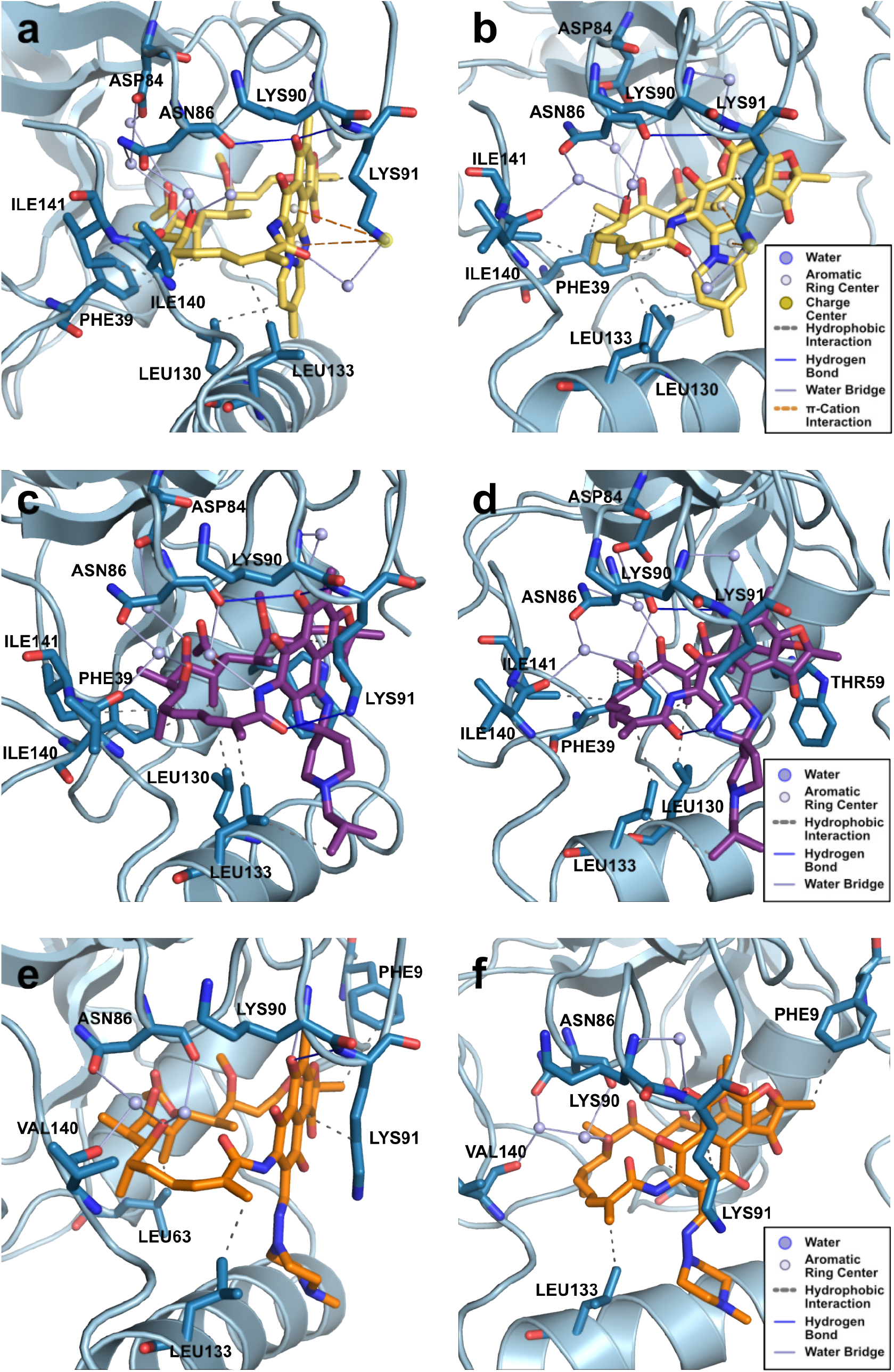
Interactions between ARR and the different rifamycins, as visualised using PLIP (Protein-Ligand Interaction Profiler) [22]. **a–b** Interactions between rifaximin and ARR. **c–d** Interactions between rifabutin and ARR. **e–f** Interactions between rifampicin and ARR.

**Supplementary Figure 3:**
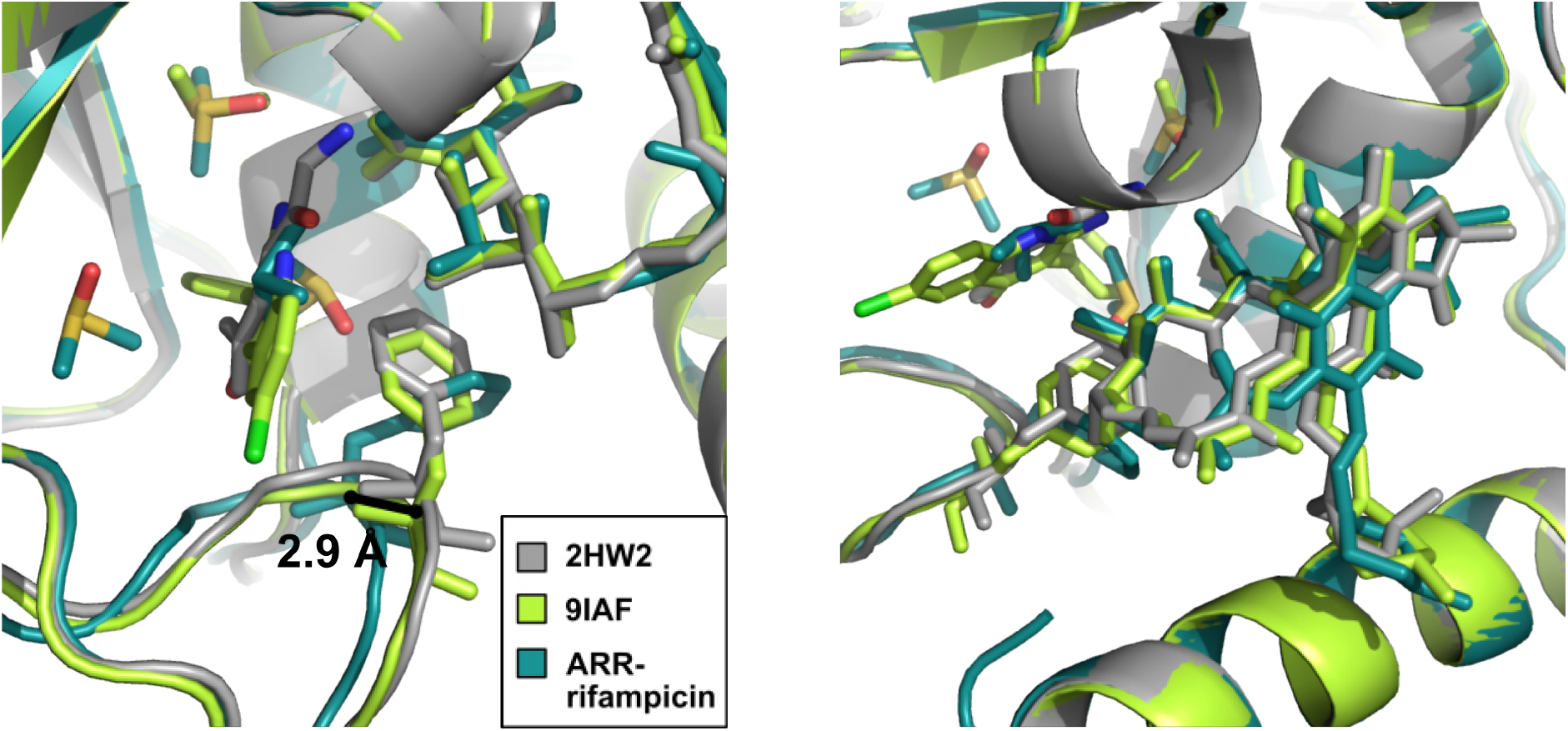
Comparison of the previously published *M. smegmatis* ARR structures in complex with rifampicin 2HW2 [10] (grey) and 9IAF [11] with our ARR-rifampicin complex.

**Supplementary Figure 4:**
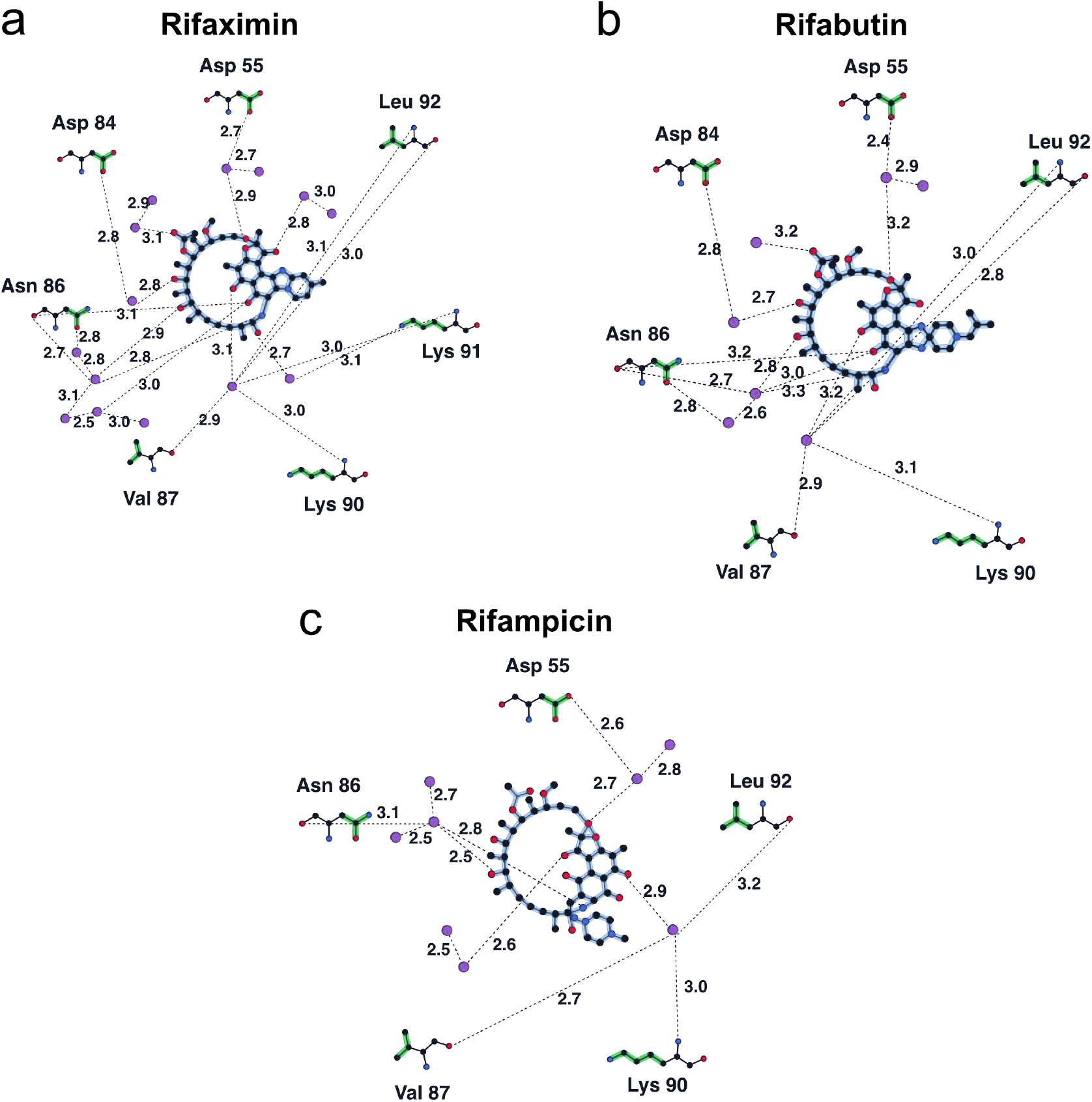
Ligand interaction plots, showing the interactions of the three different rifamycins rifaximin (**a**), rifabutin (**b**) and rifampicin (**c**) with the surrounding water molecules and amino acids.

**Supplementary Figure 5:**
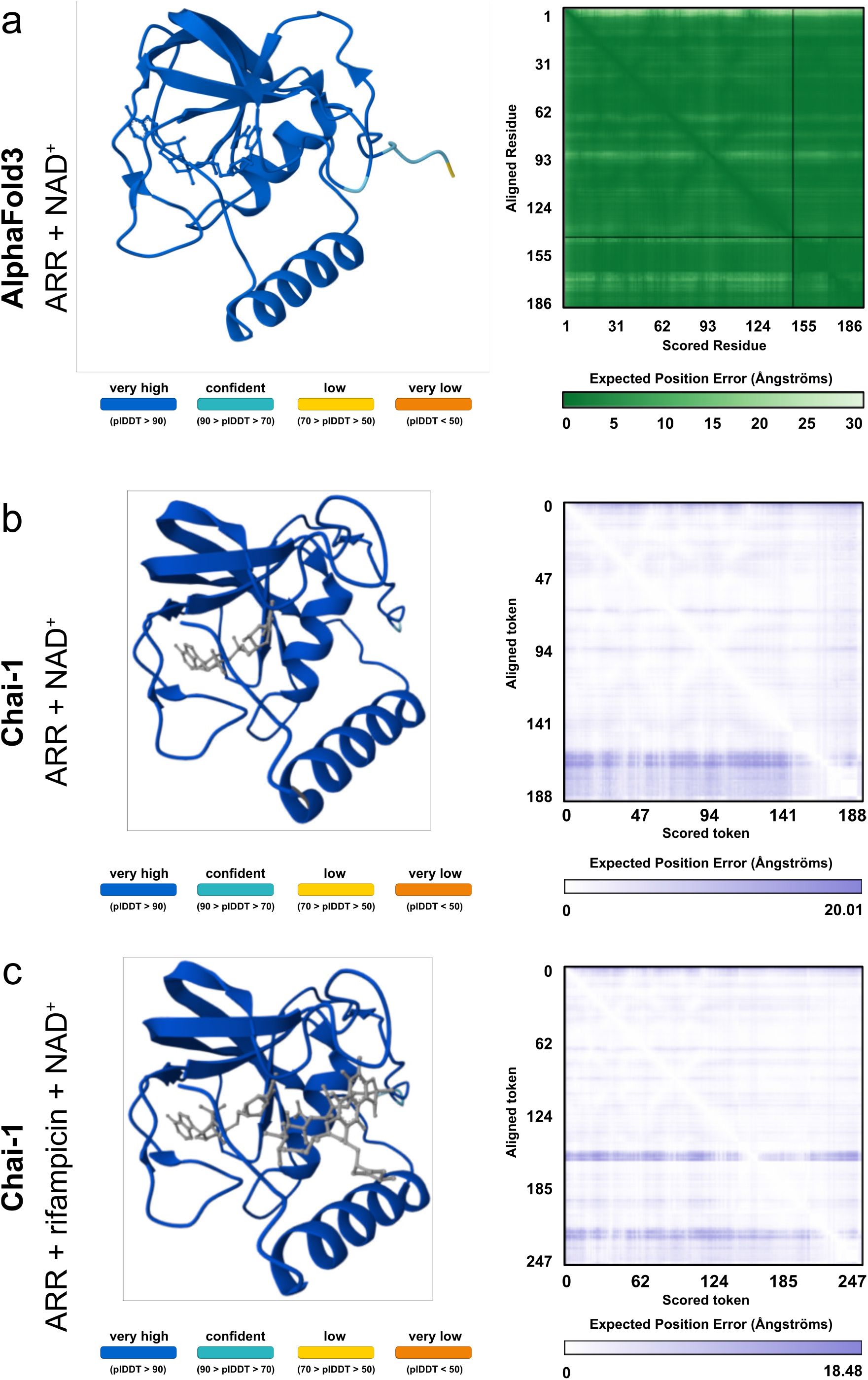
AlphaFold 3 [12] and Chai-1 [13] confidence metrics. **a** AlphaFold 3 confidence metrics for ARR + NAD^+^. **b** Chai-1 confidence metrics for ARR + NAD^+^. **c** Chai-1 confidence metrics for ARR + rifampicin + NAD^+^.

